# Danger changes the way the brain consolidates neutral information; and does so by interacting with processes involved in the encoding of that information

**DOI:** 10.1101/2022.12.02.518124

**Authors:** Omar A. Qureshi, Jessica Leake, Andrew J. Delaney, Simon Killcross, R. Frederick Westbrook, Nathan M. Holmes

## Abstract

This study examined the effect of danger on consolidation of neutral information in two regions of the rat (male and female) medial temporal lobe: the perirhinal cortex (PRh) and basolateral amygdala complex (BLA). The neutral information was the association that forms between an auditory stimulus and a visual stimulus (labelled S2 and S1) across their pairings in sensory preconditioning. We show that, when the sensory preconditioning session is followed by a shocked context exposure, the danger shifts consolidation of the S2-S1 association from the PRh to the BLA; and does so by interacting with processes involved in encoding of the S2-S1 pairings. Specifically, we show that the initial S2-S1 pairing in sensory preconditioning is encoded in the BLA and not the PRh; whereas the later S2-S1 pairings are encoded in the PRh and not the BLA. When the sensory preconditioning session is followed by a context alone exposure, the BLA-dependent trace of the early S2-S1 pairings decays and the PRh-dependent trace of the later S2-S1 pairings is consolidated in memory. However, when the sensory preconditioning session is followed by a shocked context exposure, the PRh-dependent trace of the later S2-S1 pairings is suppressed and the BLA-dependent trace of the initial S2-S1 pairing is consolidated in memory. These findings are discussed with respect to mutually inhibitory interactions between the PRh and BLA, and the way that these regions support memory in other protocols, including recognition memory in people.

**Significance Statement:** The perirhinal cortex (PRh) and basolateral amygdala complex (BLA) process the pairings of neutral auditory and visual stimuli in sensory preconditioning. The involvement of each region in this processing is determined by the novelty/familiarity of the stimuli as well as events that occur immediately after the preconditioning session. Novel stimuli are represented in the BLA; however, as these stimuli are repeatedly presented without consequence, they come to be represented in the PRh. Whether the BLA- or PRh-dependent representation is consolidated in memory depends on what happens next. When nothing of significance occurs, the PRh-dependent representation is consolidated and the BLA-dependent representation decays; but when danger is encountered, the PRh-dependent representation is inhibited and the BLA-dependent representation is selected for consolidation.

## Introduction

Animals learn about stimuli that signal motivationally significant events and use this information to guide their behavior. They also learn about the relations between stimuli that lack motivational significance, which serves as a hedge against future need. This is evident in the laboratory phenomenon of sensory preconditioning, which is studied in a three-stage protocol. In stage 1, rats are exposed to pairings of two affectively neutral stimuli, such as a sound (S2) and a light (S1) in a familiar context. In stage 2, rats are exposed to pairings of one of these stimuli, e.g., the S1, with brief but aversive foot-shock. Finally, in stage 3, rats exhibit fear responses (e.g., freezing) when tested with the S1 that had been paired with shock as well as the S2 that had never been paired with danger. Controls show that the responses to S2 are conditional on the association produced by the S2-S1 pairings in stage 1, rather than generalization from the conditioned S1 (e.g., Parkes & Westbrook, 2010; Rizley and Rescorla, 1972). Thus, while the association formed by the pairings is inconsequential to the rats in stage 1, the adaptive value of its encoding is realized at the time of testing when they treat the S2 as dangerous and respond appropriately.

Recent studies have used this protocol to examine the substrates of the S2-S1 association in two regions of the medial temporal lobe: the perirhinal cortex (PRh), which encodes information about neutral stimuli in other protocols (e.g., object recognition; (Albasser et al., 2009; Barker et al., 2006; Ennaceur et al., 1996; Mumby et al., 2002; Norman & Eacott, 2005; Warburton et al., 2005; Winters and Bussey, 2005); and the basolateral amygdala complex (BLA), which encodes information about stimuli that signal motivationally significant events (e.g., S1-shock associations; see Johansen et al., 2011; Maren, 2005; Rodrigues et al., 2004). These studies show that, when rats are exposed to S2-S1 pairings in a familiar and safe context, encoding and consolidation of the S2-S1 association typically requires neuronal activity in the PRh but not the BLA: silencing the PRh (via infusion of the GABA agonist, muscimol) immediately before or after the session of S2-S1 pairings in stage 1 significantly reduced the test level of freezing to S2; whereas silencing the BLA before or after stage 1 had no effect on this freezing (Holmes et al., 2013; Holmes et al., 2018). However, the roles of these regions are reversed when rats are exposed to a session of S2-S1 pairings in a safe context and, minutes later, shocked in that context: here, silencing the PRh immediately after the shocked context exposure in stage 1 had no effect on freezing to S2 at test; whereas silencing the BLA after the shocked context exposure in stage 1 significantly reduced this freezing (Holmes et al., 2018). That is, exposure to danger after stage 1 changes the way the brain consolidates the S2-S1 association in sensory preconditioning: it disengages the PRh and, instead, engages the BLA for this consolidation.

How does danger *after* the sensory preconditioning session shift consolidation of the S2-S1 association from the PRh to the BLA? The present study addressed this question using a combination of learning theory, pharmacology and electrophysiology. It specifically tested the hypothesis that danger in the form of the shocked context exposure has these effects because of the way that the PRh and BLA process the early and later S2-S1 pairings. The early S2-S1 pairings are processed in the BLA as their presentations are novel and consequences unknown; whereas the later S2-S1 parings are processed in the PRh as their presentations are familiar and consequences (nothing of importance) known. If nothing else happens, the BLA-dependent memory trace decays and the PRh-dependent trace is consolidated to long-term memory. If, however, animals are exposed to danger in the form of the shocked context exposure, the PRh-dependent trace is suppressed and the BLA-dependent trace is consolidated to memory.

## Materials and Methods

### Subjects

Subjects were experimentally naïve male and female Long-Evan rats, 8-12 weeks old, obtained from the Rat Breeding Facility at the University of New South Wales (Sydney, Australia). The rats were housed in plastic tubs (67 cm length × 40 cm width × 22 cm height) by sex, with a minimum of four rats per cage. These cages were kept in a colony room where the temperature was maintained at approximately 21°C, and kept on a 12-hour light/dark cycle (lights on at 0700 and off at 1900). All rats received unrestricted access to food and water for the duration of the experiment.

### Surgery and drug infusions

Prior to behavioral training, rats were surgically implanted with bilateral guide cannulas targeting the PRh or BLA. They were anesthetized with 5% of isoflurane for induction and 2-2.5% for maintenance (Cenvet, Australia) and positioned on a stereotaxic apparatus (Kopf Instruments, Tujunga, CA). Following site incision, two guide cannulas, 26-gauge, 11 mm (Plastics One, Roanoke, VA), were implanted bilaterally through holes drilled in both hemispheres of the skull. The tips of the guide cannula aimed at the PRh were: anteroposterior, -4.3 mm; mediolateral, ±5 mm; dorsoventral, -8.4 mm; angled at 9°. Those aimed at the BLA were: 2.4 mm anteroposterior; ±4.9 mm mediolateral; -8.2 mm dorsoventral (Paxinos and Watson, 2007). The cannulas were maintained in place with dental cement and four jeweler screws attached to the skull. Dummy cannulas were placed in each guide cannula and remained there at all times except during infusions. Immediately following surgery, rats received an intraperitoneal (i.p.) injection of a prophylactic (0.4 ml) dose of procaine penicillin (33 mg/kg) to prevent infection. Rats were allowed a minimum of seven days to recover, during which time they were monitored and weighed daily.

Cycloheximide, _D_-2-amino-5-phosphonopentanoic acid (DAPV) or vehicle was infused bilaterally into the BLA or PRh. On the day of infusions, the dummy cannulas were removed and 33-gauge internal cannulas inserted into the guide cannula. The internal cannulas, which projected an additional 1 mm ventral to the tip of the guide cannulas, were connected to 25 μl Hamilton syringes attached to an infusion pump (Harvard Apparatus). A total volume of 0.5 μl was infused into each hemisphere at a rate of 0.25 μl/min. Following infusion, the internal cannula remained in place for an additional two min to allow for diffusion away from the tip. One day prior to the infusions, rats were familiarized with the infusion procedure by removing their dummy cannula and turning on the infusion pump.

### Drugs

The protein synthesis inhibitor, cycloheximide, and the NMDAr antagonist, DAPV, were obtained from Sigma-Aldrich Australia. Cycloheximide was prepared in the manner described in the literature (e.g., Duvarci et al., 2005). It was first dissolved in 70% ethanol to yield a stock solution with 200 μg/μL concentration which was then diluted 1:4 with artificial cerebral spinal fluid (ACSF: Sigma-Aldrich, Australia) to a final concentration of 40 μg/μL. Vehicle was prepared by diluting 70% ethanol 1:4 with ACSF. DAPV was also prepared in the manner described previously (Duvarci et al., 2005). It was dissolved in 70% ethanol to create a stock solution of 200 μg/μl. The stock was then diluted 1:4 in ACSF to yield a final solution of 40 μg/μl. A 70% ethanol solution that was diluted 1:4 in ACSF was used as the vehicle solution. DAPV was dissolved in ACSF at a concentration of 10 μg/μL (Maren et al., 1996). Nonpyrogenic saline (0.9% w/v) was used as a vehicle solution for DAPV infusions.

### Histology

Following behavioral testing, rats were euthanized with a lethal dose of sodium pentobarbital. Their brains were removed, rapidly frozen and sectioned coronally at 40 μm through the PRh or BLA. Every second section was collected on a glass slide and stained with cresyl violet. The location of cannula tips was determined under a microscope by a trained observer, unaware of the rat’s group designation, using the boundaries defined by the atlas of Paxinos and Watson (2007). Rats with inaccurate cannula placements or with extensive damage were excluded from the statistical analysis. Figure 1 shows placement of the most ventral portion of these cannulas in the PRh and BLA for all rats that were included in the statistical analyses.

**Figure 1.**
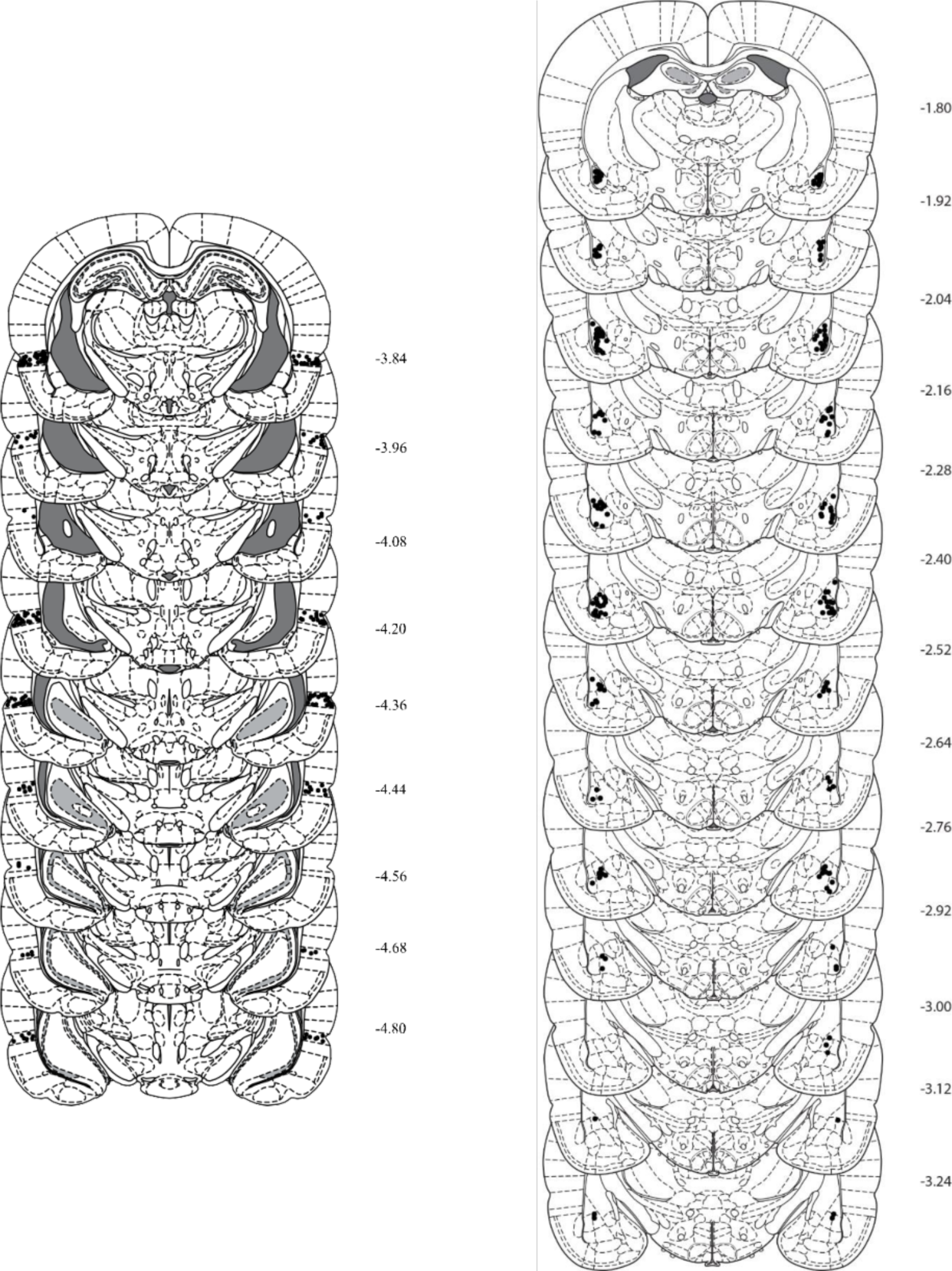
Cannula placements as verified on nissl-stained sections for rats included in experiments where we targeted the PRh (left) and BLA (right). The symbols represent the most ventral point of the cannula track for each rat on coronal sections based on the atlas of Paxinos and Watson (2007).

### Behavioral apparatus

All behavioral procedures occurred in four chambers, each measuring 30 cm (length) x 26 cm (width) x 30 cm (high). The sidewalls and ceiling of each chamber were made of aluminum while the front and back walls were made of clear Plexiglass. The floor consisted of stainless-steel rods (two mm in diameter, spaced 15 mm apart, center to center) connected to a constant-current shock generator which delivered an unscrambled AC 50 Hz shock to the floor. The shock used to render the context dangerous was 0.5 mA x 0.5 s and that used to condition S1 was 0.8 mA x 0.5 s. A tray located below the floor contained bedding material which was replaced after each rat was removed. Each chamber was enclosed in a wooden sound- and light-attenuating cabinet. The floor, ceiling and walls of the cabinet were painted black. A speaker mounted on the back wall of each cabinet delivered a 1000-Hz tone at 75 dB against a background noise of ∼45 dB, measured by a digital sound level meter (Dick Smith Electronics, Australia). A set of LEDs, also mounted on the back wall of each cabinet, delivered a light flashing at a rate of 3.5 Hz whose luminance was ∼8 lux. Each chamber was illuminated by an infrared light source (940 ± 25 nm). A camera was mounted on the back wall of each cabinet and connected to a monitor and DVD recorder located in another room in the laboratory. It was used to record the behavior of each rat. All stimulus presentations were controlled using MATLAB (MathWorks Inc.).

### Behavioral procedure

#### Context exposure

Rats received two 20 min exposures to the chambers, one in the morning and the other approximately three h later in the afternoon. These sessions were intended to familiarize the rats with the chambers and reduce any neophobic reactions.

#### S2-S1 pairings

On the next day, rats received eight S2-S1 pairings in all of the experiments apart from Experiment 2 where they received a single S2-S1 pairing. The identities of the stimuli were counterbalanced such that for half the rats, the tone and the flashing light constituted S2 and S1, respectively, and for the other half, the flashing light and the tone constituted S2 and S1, respectively. Each 30 s presentation of S2 co-terminated in the onset of a 10 s S1. The duration of the interval between each of the pairings was five min, with the first presentation of S2 occurring five min after placement in the chambers. Rats were removed from the chambers two min after the last S1 presentation and placed in holding cages. In Experiment 2, rats were exposed to the chambers for seven min, during which they received a single presentation of S2 and S1, with S2 constituting a 30 s tone and S1 was a 10 s flashing light. For those that received a paired presentation of these stimuli, S2 occurred five min after placement in the chamber and co-terminated in the onset of the 10 s S1. For those that received an unpaired presentation of these stimuli, S2 presentation occurred 150 s after placement in the chamber and S1 presentation occurred five min after placement in the chamber. Rats were removed from the chambers two min after S1 presentation and placed in holding cages.

#### Pre-extinction

Pre-extinction (Experiment 4) occurred 24 h after the S2-S1 pairings. Rats received eight S2 alone presentations. Each S2 presentation was 30 s in duration and spaced three min apart. The first presentation occurred two min after placement in the chamber and rats remained in the chambers for two min after the final presentation.

#### Additional context session

In all experiments apart from Experiment 2, rats received an additional five min exposure to the chambers five min after sensory preconditioning. In these sessions, some rats were shocked twice in the chambers: each shock was delivered at 0.5 mA for 0.5 s, the first shock occurred three min after placement in the context and the second shock occurred one min later, i.e., four min after placement in the context. The remaining rats in these experiments were not shocked.

#### S1-shock pairings

Threat conditioning occurred the day after context extinction (Experiments 1 and 3), the S2-S1 pairings (Experiment 2) or pre-extinction (Experiment 4). It consisted in four S1-shock pairings. Each presentation of the 10 s S1 co-terminated in the onset of foot-shock. The first presentation occurred five min after placement in the chambers and the interval between pairings was five min. Rats were removed from the chambers one min after the final shock and returned to their home cages.

#### Context extinction

In Experiments 1 and 3, context extinction occurred the day after the S2-S1 pairings as well as the day after threat conditioning. In the remaining experiments, context extinction occurred the day after threat conditioning. Extinction consisted in two 20 min exposures to the chambers, one in the morning and the other in the afternoon. These exposures were intended to extinguish any freezing elicited by the context and thereby provide a measure of the level of freezing to the sound and the light uncontaminated by any context-elicited freezing. Rats additionally received a further 10 min extinction exposure to the context on the day of S2 test.

#### Test

Rats were tested for their levels of freezing to S2 approximately two h after the 10 min context-extinction session. This test consisted in eight S2 alone presentations. Each presentation was 30 s in duration and spaced three min apart. The first presentation occurred two min after placement in the chamber and rats remained in the chambers for two min after the final presentation. The following day, rats were tested with S1. This test consisted in eight S1 alone presentations. Each presentation was 10 s in duration and spaced three min apart. The first S1 presentation occurred two min after placement in the chamber and rats remained in the chambers for two min after the final presentation.

### Scoring and statistics

Freezing was used as the measure of sensory preconditioned responding to S2 and conditioned responding to S1. Freezing was scored using a time sampling procedure in which each rat was observed every two s and scored as either ‘freezing’ or ‘not freezing’ by two observers, one of whom was blind to group assignment (Fanselow, 1980). The correlation between the scores of the two observers was high, > 0.9 (calculated using the Pearson correlation coefficient), and any discrepancies between the scores was resolved in favor of the blind observer. The number of observations scored as freezing was calculated and expressed as a percentage of the total number of observations.

The data were averaged across all trials and analyzed using a set of planned orthogonal contrasts (Hayes, 1963) with the type one error rate controlled at α = 0.05. Standardized 95% confidence intervals (CIs) were reported for significant results, and Cohen’s d (*d*) is reported as a measure of effect size (where 0.2, 0.5, and 0.8 is a small, medium and large effect size respectively).

### Methods for electrophysiology

#### Acute brain slice preparation

Coronal brain sections were prepared from 21-35 day old male Sprague Dawley rats (Animal Resources Centre, Perth, Western Australia) using Leica VT1000S vibratome at 0°C using previously reported techniques (Delaney et al., 2010). Brain slices were transferred to a holding chamber of low calcium/high magnesium artificial cerebro-spinal fluid (ACSF; containing (in mM) 118 NaCl, 25 NaHCO3, 10 Glucose, 0.5 CaCl2, 1.2 NaH2PO4, 2.6 MgCl2 and 0.5 Kynurenic acid) at 33-34°C for 30 mins, then maintained for several hours at room temperature. All experimental procedures were approved by the Animal Care and Ethics Committee at Charles Sturt University and performed in accordance with the National Health and Medical Research Council (Australia) *Guidelines for the Care and Use of Animals for scientific purposes* (2013).

#### Whole-cell recording

Brain slices were transferred a recording chamber continuously perfused with oxygenated ACSF solution containing (in mM): 118 NaCl, 25 NaHCO_3_, 10 Glucose, 2.5 CaCl_2_, 1.2 NaH_2_PO_4_ and 1.3 MgCl_2,_ heated to 32-33°C. Voltage-clamp whole-cell recordings were made from neurons visually identified using IR/DIC techniques using 3-3.5 MΩ patch electrodes filled with pipette solution containing (mM): CsMeSO_4_ 130, NaCl 15, HEPES 10, Mg_2_ATP 2 and Na_3_GTP 0.3 (pH 7.2 with CsOH, osmolarity 290 mOsm/kg) amplified using a Multiclamp 700B amplifier (Molecular Devices, Sunnyvale, CA). Current signals were filtered at 10 kHz and digitized at 20 kHz (National Instruments, USB-6221 digitiser), acquired, stored and analyzed on Toshiba Satellite Pro L70 using Axograph software.

During experiments, bicuculine and NBQX (Abcam) were added to the ACSF solution used to perfuse the slice, and glutamate (Sigma) was applied in a 10mM solution in ACSF, via pressure injection from a glass pipette (1-1.5 MΩ) using a 101-220 digital dispenser (Fisnar, USA). Evoked post-synaptic currents were evoked using a bipolar stimulating electrode rotated perpendicular to the slice with a single electrode tip placed onto the surface of the slice, stimulation provided by a DS2A Isolated Voltage stimulator (Digitimer, UK). Evoked current responses shown are averages of 10-20 individual trials. Puffer responses are shown as chart recordings. Access resistance (5-15 MΩ) was monitored throughout the experiments and experiments were discontinued if series resistance changed by more than 10%.

All experiments are within-subject experiments where a subject is a typical PRh layer 5 pyramidal neuron or an LA pyramidal neuron. In each experiment a single treatment is tested using a single measure (e.g., current amplitude) except where a washout and second treatment was specified. Each replicate in an experiment is conducted by recording from single neurons in separate brain slices and the identity of the neuron was established visually and using cellular properties (e.g., input resistance). Within each experiment brain slices are prepared from at least two rats. Paired *t-*tests performed in Microsoft Excel were used for statistical comparisons between measures in control and in treatment (except where indicated). All results are expressed as mean ± SEM and percentage change is calculated as absolute percentage change.

## Results

### Experiments 1A and 1B: Exposure to danger after sensory preconditioning disengages the PRh and engages the BLA for consolidation of the S2-S1 association

The aim of Experiments 1A and 1B was to confirm that exposure to danger after sensory preconditioning (in the form of a shocked context exposure) disengages the PRh for consolidation of the S2-S1 association, and instead, engages the BLA (Holmes et al., 2018). In each experiment, four groups of rats were surgically prepared with cannulas targeting either the PRh (Experiment 1A) or BLA (Experiment 1B). Following recovery from surgery, all rats were familiarized with the conditioning chambers and then exposed to eight 30 s presentations of S2, each of which terminated in presentation of a 10 s S1. The identity of S2 and S1 (auditory and visual) was counterbalanced. Following the final S2-S1 pairing, rats were removed for a few minutes and returned to the chambers where rats in two groups were shocked and those in the other two were not shocked. One group in each pair then received a PRh (Experiment 1A) or BLA (Experiment 1B) infusion of the protein synthesis inhibitor, cycloheximide (CHX), as protein synthesis has been shown to be critical for the cellular changes that consolidate new information in memory (Hernandez & Abel, 2008; Johansen et al., 2011; Lay et al., 2018; Leidl et al., 2018; Rodrigues et al., 2004; Williams-Spooner et al., 2019). The other group in each pair received an infusion of vehicle (VEH) into the PRh or BLA. There were thus four groups in each experiment, labelled Groups Shock-CHX, No Shock-CHX, Shock-VEH and No Shock-VEH, with the experiments differing according to the region targeted for the infusion. All rats were subsequently exposed to S1-shock pairings and finally tested first with S2 and then with S1.

In Experiment 1A, the test levels of freezing to S2 or S1 did not significantly differ between rats in Groups Shock-VEH and No Shock-VEH (*F* < 1). Therefore, these groups were combined into a composite vehicle-infused control group (Group VEH). This was also the case in Experiment 1B and all subsequent experiments. In Experiment 1A, baseline levels of freezing in the two test sessions were low (< 10%) and equivalent in each of the groups (*F*s < 1). This was also the case in Experiment 1B (*F*s < 1) and for all subsequent experiments (data not shown).

Conditioning was successful in both experiments. In Experiment 1A, freezing to S1 increased across its pairings with shock, *F*_1,32_ = 99.308, p < 0.01, 95% CI [1.257, 1.903]. The rate of increase did not differ between rats in Groups Vehicle and Shock-CHX, *F* < 1. However, it did differ between rats in these two groups and those in Group No Shock-CHX, *F*_1,32_ = 6.897, p < 0.05, 95% CI [0.197, 1.545]. This difference was unexpected, but importantly, there were no significant differences between the three groups in freezing to S1 on its final pairing with shock, *F*s < 4, or in the overall levels of freezing across the four pairings, *F*s < 1.8. In Experiment 1B, freezing to S1 also increased across its four pairings with shock, *F*_1,29_ = 68.216, p < 0.05, 95% CI = [1.356, 2.248]. There were no significant differences in the rate at which freezing increased between the groups, *F*s < 1, and no significant between-group differences in the overall levels of freezing across the pairings, *Fs* < 2.6.

Figure 2 shows the mean (+SEM) levels of freezing to S2 (left) and S1 (right) during test sessions in Experiments 1A (Figure 2B, 2C) and 1B (Figure 2E, 2F). Inspection of Figure 2B suggests that sensory preconditioning of S2 required protein synthesis in the PRh when the session of S2-S1 pairings was followed by context alone exposure (Group No Shock-CHX) but *not* when this session was followed by a shocked exposure to the context (Figure 2C); conversely, sensory preconditioning did not require protein synthesis in the BLA when the session of S2-S1 pairings was followed by context alone exposure (Group No Shock-CHX) but *did* require protein synthesis when the session was followed by a shocked exposure to the context (Figure 2B). Inspection of Figure 2F suggest that first-order conditioning of S1 was similar in all of the groups in each experiment.

**Figure 2.**
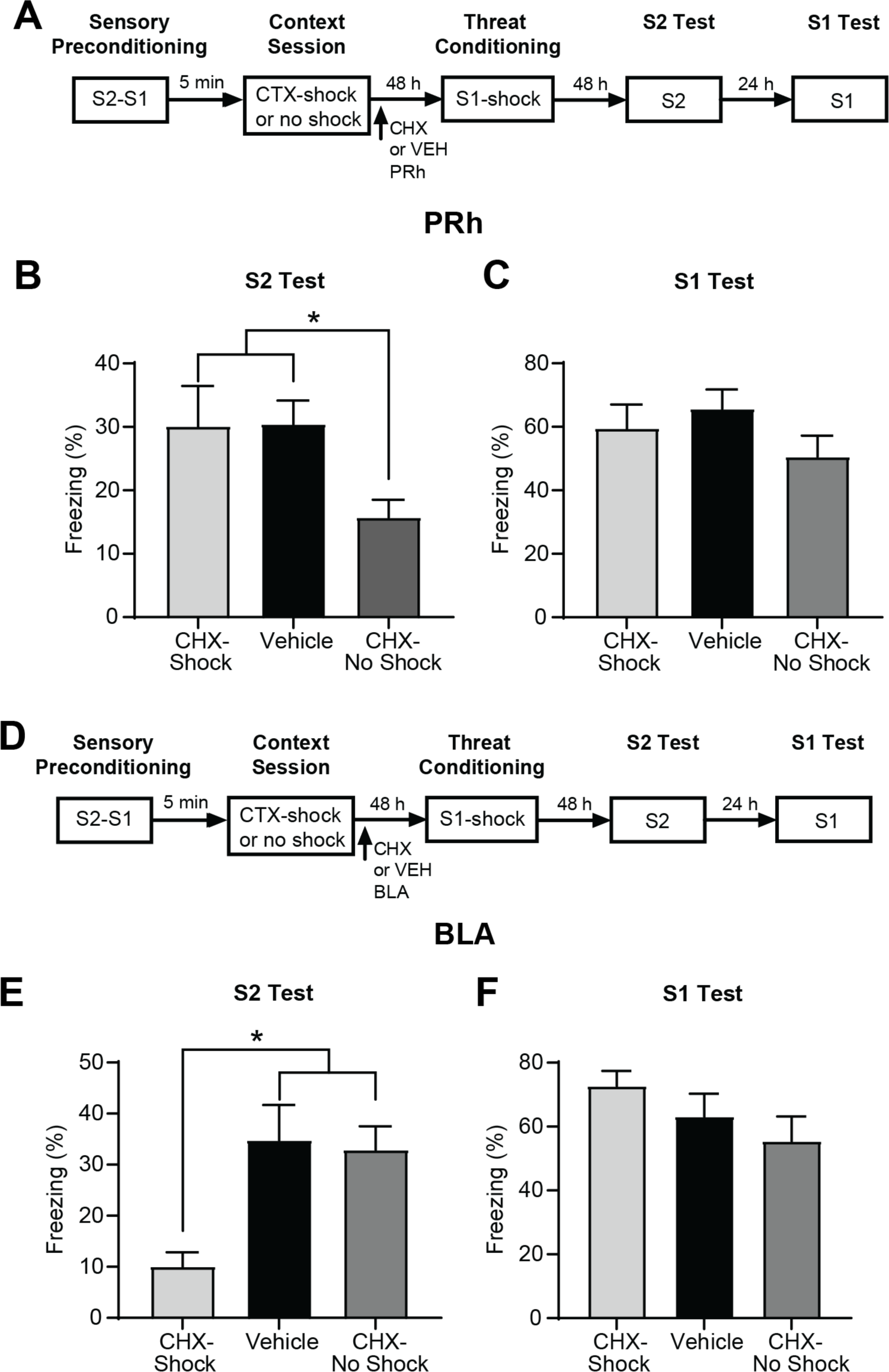
Design of Experiments 1A (A) and 1B (D). Mean (+SEM) levels of freezing on the S2 and S1 tests in Experiments 1A (B-C) and 1B (E-F). The freezing levels have been averaged over the eight stimulus presentations that occurred in each test session. A cycloheximide infusion into the PRh disrupted sensory preconditioned fear of S2 in rats that were not shocked (Group CHX-No Shock) but not in rats that were shocked (Group CHX-Shock (B). A cycloheximide infusion into the BLA disrupted sensory preconditioned fear of S2 in rats that were shocked (Group CHX-Shock) but not in rats that were not shocked (Group CHX-No Shock) (E). There were no effects of the PRh or BLA cycloheximide infusion on conditioned fear of S1 (C, F).

These impressions were supported by the statistical analysis. In Experiment 1A, rats in Group No Shock-CHX froze significantly less to S2 than those in Groups Vehicle and Shock-CHX, *F*_1,32_ = 8.179, p < 0.01, 95% CI = [0.288, 1.716], who did not differ from each other, *F* < 1. In contrast, in Experiment 1B, rats in Group Shock-CHX froze significantly less to S2 than those in Groups Vehicle and No-Shock-CHX, *F*_1,32_ = 13.200, p < 0.01, 95% CI = [0.581, 2.065], who did not differ from each other, *F* < 0.1. Finally, in both experiments, conditioning of S1 was unaffected by whether or not rats had been shocked after the session containing the S2-S1 pairings and/or infused with cycloheximide or vehicle; there were no significant differences between the groups in freezing to the S1, (*F*s < 2.3 and < 2.5 in Experiments 1A and 1B, respectively). Thus, the differences in freezing to S2 cannot be attributed to effects of the manipulations on freezing *per se*. Instead, the results of Experiments 1A and 1B are consistent with the proposal that exposure to danger shifts consolidation of the S2-S1 association from the PRh to the BLA: hence, the PRh cycloheximide infusion disrupted freezing to S2 among rats that had not been shocked but failed to do so among rats that had been shocked; and the BLA cycloheximide infusion disrupted freezing to S2 among rats that had been shocked but not among rats that had not been shocked.

### Experiments 2A and 2B: The initial S2-S1 pairing is encoded in the BLA and not the PRh

How can danger *after* the session containing the S2-S1 pairings cancel the involvement of the PRh in consolidation and instead recruit the BLA? We propose that this effect of danger reflects the way that the S2-S1 association is encoded. The idea is that the initial S2-S1 pairings are in fact processed in the BLA as each of the stimuli, although affectively neutral, are novel and the consequences of the S2-S1 episodes are unknown. In contrast, the later S2-S1 pairings are processed in the PRh as the stimuli have become familiar and the consequences of the episodes known (nothing or nothing of importance). What occurs shortly after the session containing the pairings then determines which of the S2-S1 traces is consolidated in long-term memory. When this session is followed by re-exposure to the familiar and safe context, the BLA-dependent trace decays and the PRh-dependent trace is consolidated in the PRh; but when the session is followed by a shocked re-exposure to the context, the PRh-dependent trace is suppressed and the BLA-dependent trace is consolidated in long-term memory (Holmes et al., 2018).

A critical prediction of this dual trace proposal is that the initial S2-S1 pairing is encoded in the BLA and not the PRh. Experiments 2A and 2B tested this prediction. Specifically, Experiment 2A examined whether a single S2-S1 pairing was sufficient to produce sensory preconditioned fear of S2; while Experiment 2B examined whether the BLA or PRh encoded the association produced by that single S2-S1 pairing. In Experiment 2A, two groups of rats were exposed to a single presentation of S2 and a single presentation of S1 in stage 1. For rats in Group P, S2 and S1 were paired such that the presentation of S2 terminated in the onset of S1; while for those in Group U, the stimuli were presented apart such that the presentation of S2 was followed several minutes later by S1. Rats in both groups were then exposed to S1-shock pairings in stage 2 and, finally, tested with S2 and S1 in stage 3. In Experiment 2B, three groups of rats underwent surgery for implantation of cannulas in the BLA or PRh and, following recovery, were trained using the protocol described for Group P. Immediately prior to the session containing the single S2-S1 pairing in stage 1, rats in one group received an infusion of the NMDA receptor (NMDAR) antagonist, DAPV, into the PRh (Group PRh-DAPV); rats in a second group received a DAPV infusion into the BLA (Group BLA-DAPV); and, finally, rats in a third group received an infusion of vehicle into the PRh or BLA (equal numbers for a composite Group Control). The NMDAR antagonist, DAPV, was selected for use here as its infusion into the PRh, but not the BLA, disrupts formation of the association produced by multiple S2-S1 pairings in a familiar, safe context whereas its infusion into the BLA, but not the PRh, disrupts formation of the association produced by multiple S2-S1 pairings in an already dangerous context (Holmes et al., 2013). The question of interest was whether encoding of the association produced by a **single** S2-S1 pairing in a familiar safe context requires NMDAR activation in the BLA and not the PRh.

Conditioning was successful. In both experiments, freezing to S1 increased across its pairings with shock (Exp 2A: *F*_1,6_ = 28.992, p < 0.05, 95% CI = [0.849, 2.267]; Exp 2B, *F*_1,20_ = 347.594, p < 0.01, 95% CI [2.297, 2.876]), and did so at an equivalent rate among the groups, *F*s < 2.263. There was no statistically significant difference in overall freezing to S1 between Groups P and U in Experiment 2A, *F* < 1. There was, however, an unexpected between-group difference in overall freezing to S1 in Experiment 2B: rats in Group PRh-DAPV exhibited significantly less freezing than rats in Group Control, *F*_1,20_ =13.898, p < 0.05, 95% CI [0.444, 1.573]. This difference notwithstanding, the two groups displayed equivalent freezing to S1 on the final conditioning trial of stage 1, *F* < 2.530; and there were no statistically significant differences in overall freezing to S1 between the other groups, *F*s < 1.

Figure 3 shows the mean (+SEM) levels of freezing to S2 (left) and S1 (right) during the test sessions in Experiments 2A (Figure 3B, 3C) and 2B (Figure 3E, 3F). Experiment 2A revealed that a single S2-S1 pairing is sufficient to produce sensory preconditioned freezing to S2: rats in Group P froze significantly more to S2 than those in Group U, *F*_1,6_ =16.330, p < .05, 95% CI = [1.127, 4.588]. Experiment 2B revealed that encoding of the association produced by the single S2-S1 pairing requires NMDAR activation in the BLA and not the PRh: rats in Group BLA DAPV froze significantly less than the weighted average of freezing by rats in Groups Vehicle and PRh DAPV, *F*_1,20_ = 4.550, p < 0.05, 95% CI [0.021, 1.860], who did not differ from each other in their levels of freezing, *F* < 1.1. Finally, in both experiments, there were no significant between-group differences in freezing to S1, *F*s < 1.414. Thus, in each experiment, the differences in freezing to S2 cannot be attributed to differences in conditioning to the S1. Instead, the results show that a single pairing of the novel S2 and S1 is sufficient to produce the association that underlies sensory preconditioned fear of S2. The results also show that the association produced by that pairing requires NMDAR activation in the BLA and not the PRh, consistent with the proposal that the initial S2-S1 pairing is encoded in the BLA and not the PRh.

**Figure 3.**
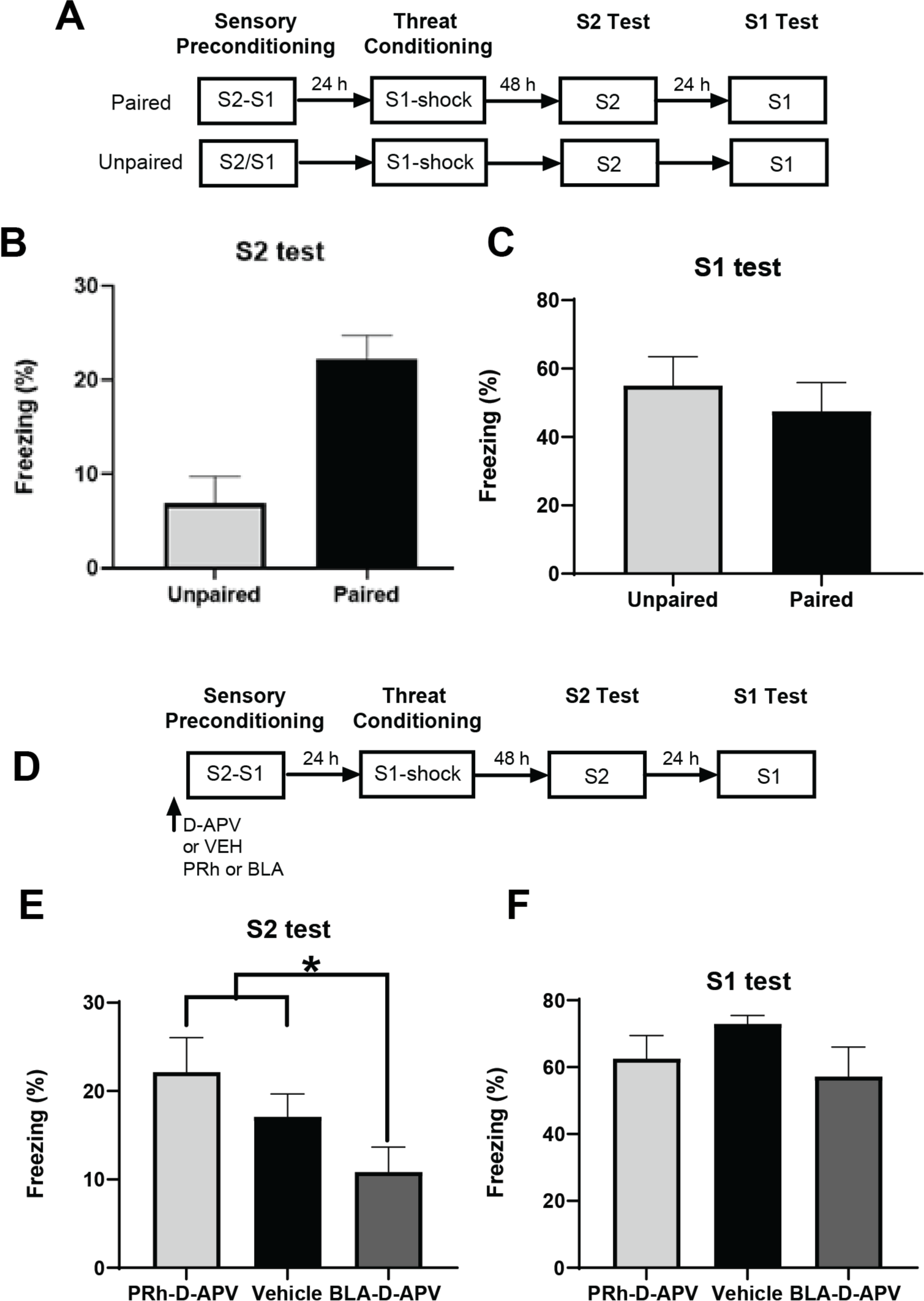
Experimental design for Experiment 2A (A) and 2B (D). Mean (+SEM) levels of freezing to S2 and S1 at test in Experiments 2A (B-C) and 2B (E-F). The freezing levels have been averaged over the eight stimulus presentations that occurred in each test session. (B-C) A single S2-S1 pairing is sufficient to produce sensory preconditioning, as rats that received this single pairing in stage 1 froze more to S2 at test than did rats that had been exposed to explicitly unpaired presentations of the S2 and S1. (E-F) The association produced by the single pairing requires neuronal activity in the BLA and not the PRh, as rats that received an infusion of the NMDAR antagonist, DAPV into the BLA prior to the single S2-S1 pairing froze less to the S2 at test than did rats that had received an infusion of DAPV into the PRh and vehicle-infused controls.

### Experiments 3A and 3B: The presence or absence of danger shortly after sensory preconditioning determines whether the PRh or the BLA is required for consolidation of the S2-S1 association

The dual trace proposal holds that, when rats are exposed to multiple S2-S1 pairings, the association is initially encoded in the BLA as the stimuli are novel and their consequences unknown, subsequently encoded in the PRh as the stimuli become familiar, and that events after the preconditioning session determine which of the traces is consolidated to long-term memory. When the session containing these pairings is followed by nothing important (such as re-exposure to the familiar, safe context), the BLA-dependent trace decays and the PRh-dependent trace is consolidated in long-term memory; however, when the session is followed by danger (such as a shocked re-exposure to the context), the PRh-dependent trace is suppressed and the BLA-dependent trace is consolidated in long-term memory.

The previous experiments confirmed a critical prediction of this proposal: that the initial S2-S1 pairing is encoded in the BLA and not the PRh. The present experiments tested two further predictions. The first is that a shocked re-exposure to the chambers immediately after the S2-S1 pairings effectively cancels the involvement of the PRh in sensory preconditioning. The second is that the shocked re-exposure engages the BLA for consolidation of the S2-S1 association but only if the BLA had been available across the S2-S1 pairings. More specifically, as long as the BLA encodes the early/novel S2-S1 pairings in sensory preconditioning, it can be engaged to consolidate the S2-S1 association when the preconditioning session is followed by danger. If, however, the BLA is not available to encode the initial S2-S1 pairing, it *cannot* be engaged to consolidate the association when the session is followed by danger: under such circumstances, danger suppresses the PRh and, thereby, disrupts sensory preconditioning.

Experiments 3A and 3B tested these predictions. In each experiment, four groups of rats were surgically prepared with cannulas targeting the PRh (Experiment 3A) or BLA (Experiment 3B). Following recovery from surgery, all rats were trained in the sensory preconditioning protocol used in Experiment 1, which commenced with a session of eight S2-S1 pairings (stage 1). The rats differed in their treatment before and after this session. Before the session, rats in two groups received an infusion of the NMDAR antagonist, DAPV, while rats in the remaining two groups received an infusion of vehicle only. After the session, rats in one group in each pair were re-exposed to the context and shocked (Groups DAPV-Shock and VEH-shock) while those in the other group of each pair were re-exposed to the context but not shocked (Groups DAPV-No shock and VEH-No shock). All rats were then exposed to S1-shock pairings in stage 2 and, finally, tested with S2 and S1 in stage 3.

As noted previously, in each experiment, rats in Groups VEH-Shock and VEH-No shock displayed equivalent levels of freezing across conditioning and testing; hence, rats in these groups were combined to form a single composite control group. In both experiments, the dual trace proposal predicts that the test level of freezing to S2 would be determined by whether or not rats were exposed to the S2-S1 pairings under a DAPV infusion and whether or not they were then re-exposed to the context and shocked or not shocked. Specifically, in Experiment 3A, a PRh infusion of DAPV would impair sensory preconditioning when the session of S2-S1 pairings was followed by a non-shocked re-exposure to the context (Group DAPV-No shock). In contrast, this infusion would spare sensory preconditioning when the session was followed by the shocked exposure to the context (Group DAPV-Shock) because that experience cancels the involvement of the PRh and, instead, engages the BLA. In Experiment 3B, we expected that a BLA infusion of DAPV would spare sensory preconditioning when the session of S2-S1 pairings was followed by a non-shocked re-exposure to the context. In contrast, this infusion would impair sensory preconditioning when the session was followed by the shocked re-exposure to the context because that experience cancels the involvement of the PRh in consolidation in spite of it having encoded the S2-S1 association. Thus, the specific predictions were: in Experiment 3A, where the target region was the PRh, rats in Group DAPV-No shock would freeze less when tested with S2 than those in Groups Control and DAPV-Shock; in Experiment 3B, where the target region was the BLA, rats in Group DAPV-Shock would freeze less when tested with S2 than those in Groups Control and DAPV-No shock.

Conditioning of S1 was successful in both experiments. Freezing to S1 increased across its four pairings with shock in Experiments 3A, *F*_1,32_ = 130.257, p < 0.05, 95% CI [0.761, 1.091] and 3B, *F*_1,28_ = 74.871, p < 0.01, 95% CI [1.403, 2.273]. There were no differences in the rate of increase amongst the groups in either experiment, *F*s < 1.461; and there were no statistically significant between-groups differences in the overall levels of freezing, *F*s < 2.738.

Figure 4 shows the mean (+SEM) levels of freezing to S2 (Figure 4B, 4C) and S1 (Figure 4C, 4F) at test in Experiments 3A and 3B. Experiment 3A revealed that NMDAR activity in the PRh was required to encode the S2-S1 association, but only among rats that were not shocked when re-exposed to the context after the S2-S1 pairings. Specifically, rats in Group DAPV-No shock froze significantly less to S2 than those in Groups Vehicle and DAPV-Shock, *F*_1,28_ = 9.778, p < 0.05, 95% CI [0.406, 1.946], who did not differ from each other, *F* < 1.974. By contrast, Experiment 3B revealed that NMDAR activity in the BLA was required to encode the S2-S1 association, but only among rats that were shocked when re-exposed to the context after the S2-S1 pairings. Specifically, rats in Group DAPV-Shock froze significantly less to S2 than those in Groups Vehicle and DAPV-No shock, *F*_(1,32)_ = 5.905, p < 0.05, 95% CI [0.136, 1.548], which did not differ from each other, *F* < 1.669.

**Figure 4.**
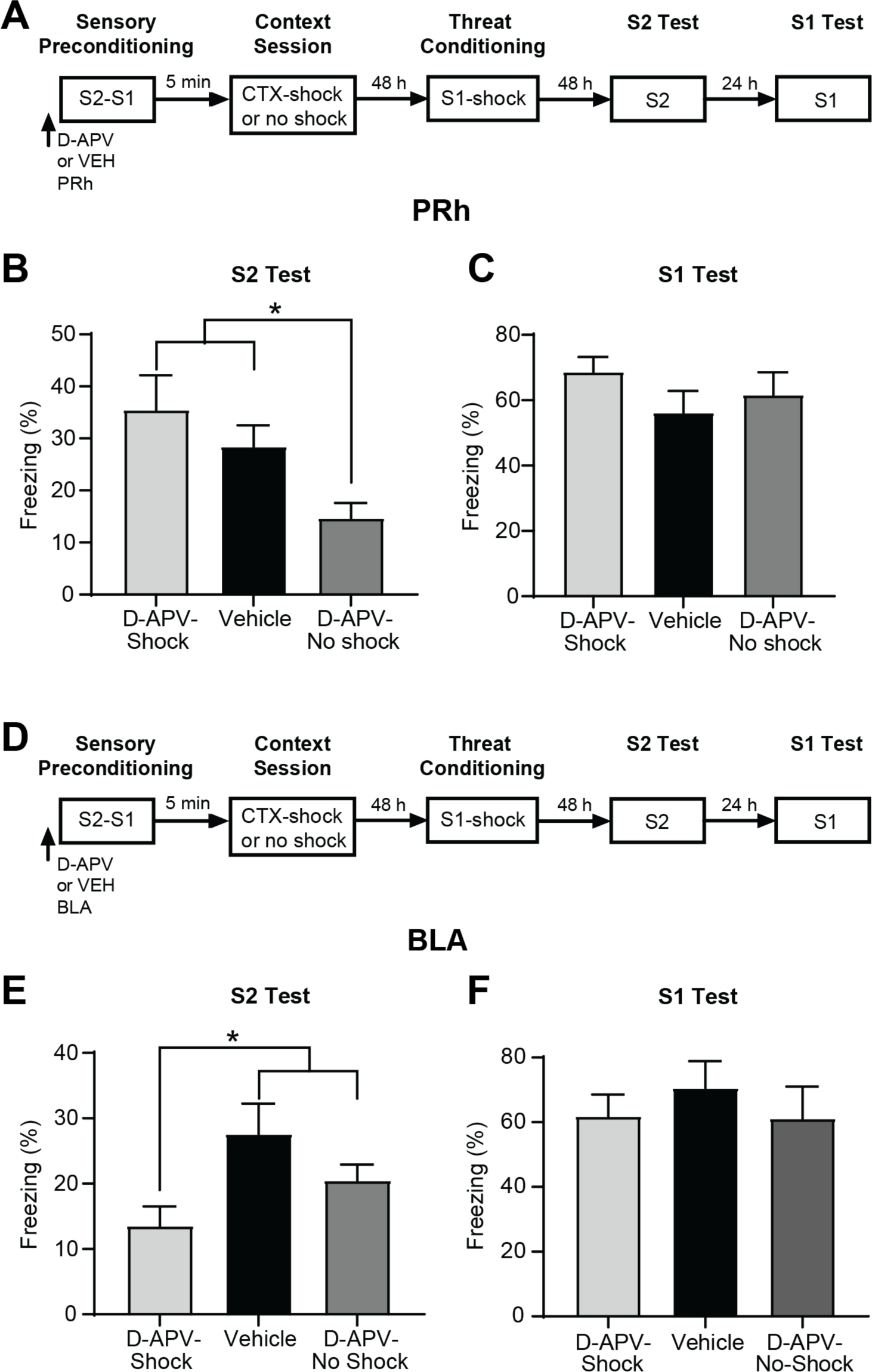
Design of Experiments 3A (A) and 3B (D). Mean (+SEM) levels of freezing to S2 and S1 during the test sessions in Experiments 3A (B-C) and 3B (E-F). The freezing levels have been averaged over the eight stimulus presentations in each test session. (B-C) Exposure to danger after the sensory preconditioning session cancels the involvement of the PRh in consolidating the S2-S1 association. (E-F) Exposure to danger after the sensory preconditioning session engages the BLA for consolidating the S2-S1 association, but only if this region was available across that session.

Finally, in both experiments, conditioning of S1 was unaffected by a DAPV infusion or foot shock in stage 1 as there were no statistically significant, between-group differences in freezing levels to S1 (*F*s < 1.975). Thus, as noted previously, the differences in freezing to S2 at test cannot be attributed to differences in the level of conditioning to S1. Instead, this experiment provides support for the dual trace proposal by showing that the presence or absence of danger shortly after the session containing the S2-S1 pairings determines the involvement of both the PRh and/or BLA in consolidating the association produced by the pairings. In the absence of danger, this association is consolidated in the PRh and not the BLA. In the presence of danger, the involvement of the PRh is suppressed and the association is consolidated in the BLA.

### Experiments 4A and 4B. Danger after sensory preconditioning changes the way that the PRh and BLA process subsequent information about the preconditioned stimulus

We next examined whether the shocked re-exposure to the context where the S2-S1 pairings had just occurred affects the learning produced by subsequent presentations of the S2 in the absence of the associated S1, so-called pre-extinction (Coppock, 1958). Rats exposed to such S2 alone presentations exhibit little or no sensory preconditioned freezing when tested with S2 (Parkes and Westbrook, 2010; Holmes et al., 2013). We have previously shown that pre-extinction of the S2-S1 association that had been formed and consolidated in a familiar, safe context requires activation of NMDAR in the PRh and not the BLA, whereas pre-extinction of the association that had been formed and consolidated in an already dangerous context requires NMDAR in the BLA and not the PRh (Holmes et al., 2013). Here we exposed rats to S2-S1 pairings in a familiar, safe context but then shocked them in that context to assess whether danger *after* the pairings alters the substrates underlying subsequent pre-extinction. Given the results of the previous experiment showing that danger suppresses the PRh and engages the BLA for consolidation of the S2-S1 association, we predicted that pre-extinction would cease to require activation of NMDAR in the PRh when the session containing the S2-S1 pairings is followed by danger and, instead, require activation of NMDAR in the BLA.

Experiments 4A and 4B tested these predictions. In each experiment, four groups of rats were surgically prepared with cannulas targeting the PRh (Experiment 4A) or BLA (Experiment 4B). Following recovery from surgery, all rats were exposed to S2-S1 pairings and, two days later, to S2 alone presentations. The rats differed in their treatment immediately after the session containing the S2-S1 pairings: those in two groups were returned to the context and shocked (Groups Shock) while rats in the remaining two groups were returned but not shocked (Groups No shock). Two days later, one group in each of these pairs received an infusion of the NMDAR antagonist, DAPV immediately prior to the session of S2 alone exposures (Groups Shock-DAPV and No shock-DAPV); while rats in the other group in each pair received an infusion of vehicle (Groups Shock-VEH and No shock-VEH). All rats were then exposed to S1-shock pairings in stage 3 and, finally, tested with S2 and then with S1 in stage 4.

As before, rats in the two vehicle-infused groups in each experiment exhibited equivalent levels of freezing across conditioning and testing. Hence, they were combined into a single, composite control (Group VEH), thereby reducing the design to three groups in each experiment. We expected to replicate our previous finding that pre-extinction requires activation of NMDAR in the PRh, not the BLA when the original S2-S1 association was formed and consolidated in a familiar, safe context (Holmes et al., 2013). That is, in the experiment which targeted the PRh (4A), pre-extinction will be impaired among rats in Group No shock-DAPV, as would be indicated by these rats freezing more when tested with S2 than rats in Group Shock-DAPV and Group VEH. We additionally expected that, when rats were shocked upon re-exposure to the hitherto safe context where the S2-S1 association had just been formed, pre-extinction will cease to require activation of NMDAR in the PRh and, instead, requires activation of NMDAR in the BLA. That is, in Experiment 4B (BLA), rats in Group Shock-DAPV would be impaired in extinguishing the S2-S1 association, as would be indicated by these rats freezing more when tested with S2 than rats in Group No Shock-DAPV and Group VEH.

Conditioning of S1 was successful in both experiments. Freezing to S1 increased across its four pairings with shock in Experiments 4A, *F*_1,32_ = 113.319, p < 0.01, 95% CI [1.391, 2.049] and 4B, *F*_1,26_ = 16.305, p < 0.01, 95% CI [0.465, 1.430]. There were no statistically significant differences in the rate at which freezing increased amongst the groups, *F*s < 1, or in their overall levels of freezing in each experiment, *F*s < 1.

Figure 5 shows the mean (+SEM) test levels of freezing to S2 and S1 in Experiments 4A (Figure 5B, 5C) and 4B (Figure 5E, 5F). Inspection of Figures 5B-C suggests that pre-extinction of the S2-S1 association requires activation of NMDAR in the PRh but only among rats that had *not* been shocked immediately after the session containing the S2-S1 pairings. The statistical analysis confirmed this suggestion: it showed that rats in Group DAPV-No Shock froze significantly more to S2 than those in Groups Composite Vehicle and DAPV-Shock, *F*_1,32_ = 8.383, p < 0.01, 95% CI [0.321, 1.846], who did not differ from each other in their levels of freezing, *F* < 1. By contrast, Figure 5E suggests that pre-extinction of the S2-S1 association requires activation of NMDAR in the BLA but only among the rats that *had* been shocked after the session containing the S2-S1 pairings. The analysis again confirmed this suggestion: it showed that rats in Group DAPV-Shock froze significantly more to S2 than those in Groups Composite Vehicle and DAPV-No Shock, *F*_1,26_ = 5.325, p < 0.03, 95% CI [0.099, 1.705], who did not differ from each other in their levels of freezing, *F* < 1.23. Finally, in both experiments, conditioning of S1 was unaffected by the shock administered after stage 1 or the drug infusion before stage 2 as there were no significant between-group differences in freezing to S1, (*F*s < 3.52 in Experiment 4A and *F*s < 0.60 in Experiment 4B). Thus, the differences in freezing to S2 in each experiment cannot be due to differences in the level of conditioning to S1. Instead, the differences in freezing to S2 show that danger after sensory preconditioning determines the substrates involved in future learning about the S2. When rats had not been exposed to danger after the sensory preconditioning session, pre-extinction of the S2-S1 association across a series of S2 alone exposures required activation of NMDAR in the PRh and not the BLA. When rats had been exposed to danger after sensory preconditioning, pre-extinction of the S2-S1 association required activation of NMDAR in the BLA and not the PRh.

**Figure 5.**
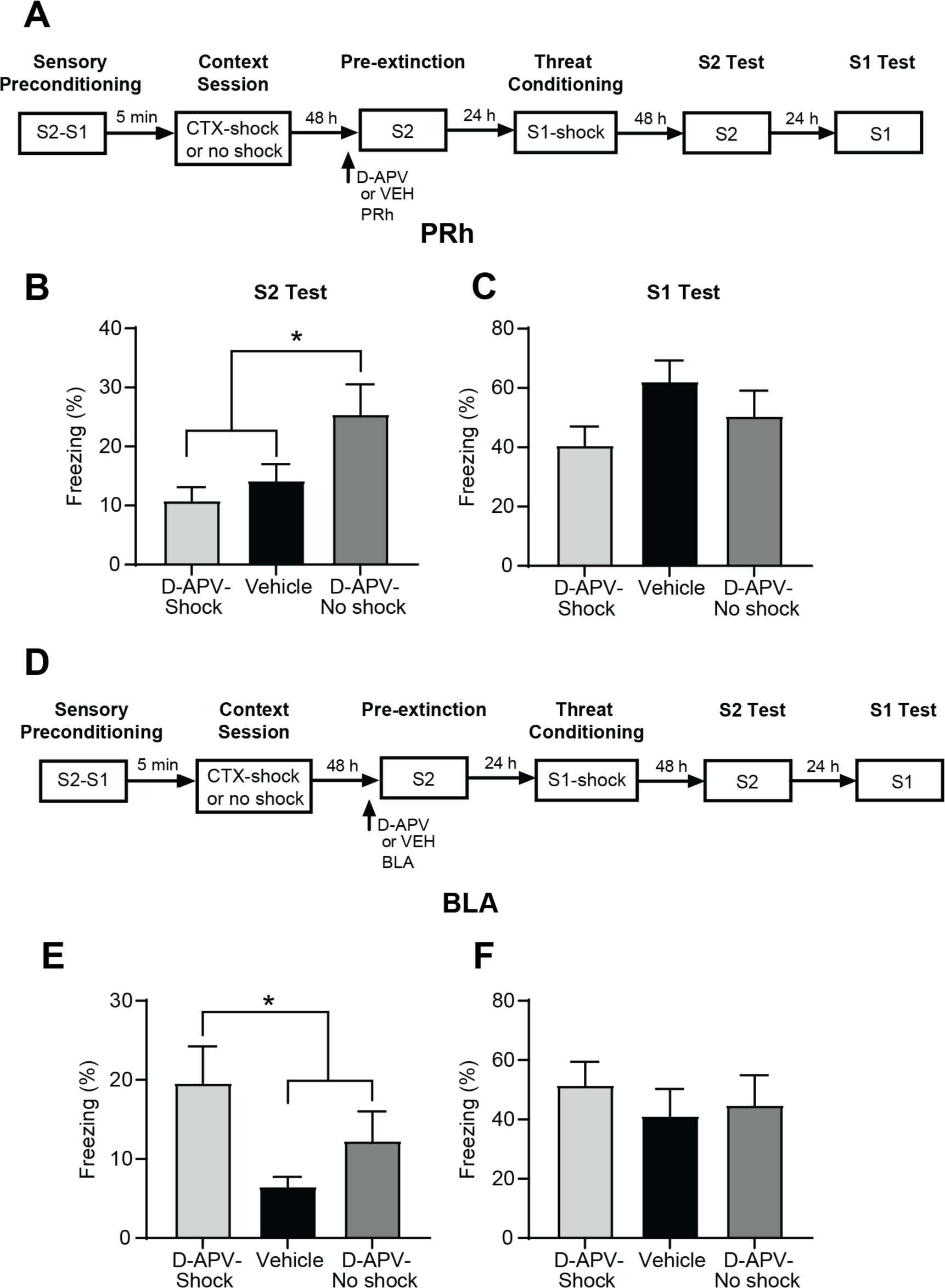
Design for Experiments 4A (A) and 4B (D). Mean (+SEM) test levels of freezing to S2 and S1 in Experiments 4A (B-C) and 4B (E-F). The freezing levels have been averaged over the eight stimulus presentations in each test session. (B-C) Exposure to danger after the sensory preconditioning session cancels the involvement of the PRh in pre-extinction of the S2-S1 association. (E-F) Exposure to danger after the preconditioning session engages the BLA for pre-extinction of the S2-S1 association.

### Experiments 5A and 5B. How do the PRh and BLA compete for processing of the neutral S2-S1 association in sensory preconditioning?

The PRh and BLA have been shown to cooperate during aversive conditioning of complex stimuli in rats (Kholodar-Smith et al., 2008), appetitive trace conditioning in cats (Paz et al., 2006), object recognition memory in rats (Roozendaal et al., 2008) and memory for emotive/neutral pictures in people (Ritchey et al., 2019). By contrast, our findings suggest that the PRh and BLA *compete* for associative formation in stage 1 of sensory preconditioning. The BLA processes the novel S2-S1 pairings and encodes the resultant S2-S1 association while the PRh is not engaged for this processing; however, as the pairings are repeated and stimuli become familiar, the PRh is engaged to encode the S2-S1 association and the BLA is disengaged. Accordingly, we next examined the potential for mutually inhibitory interactions between the BLA and PRh: specifically, whether: 1) increased activity in the BLA results in functional inhibition of the PRh; and 2) increased activity in the PRh results in functional inhibition of the BLA. To do so, we used slice electrophysiology and stimulated: 1) the lateral region of the BLA (LA) while recording from pyramidal neurons in the PRh; and 2) the PRh while recording from pyramidal neurons in the LA. We expected that stimulating activity in one region would increase inhibitory post-synaptic currents in the other. Such findings would indicate capacity for the BLA and PRh to mutually inhibit each other during stage 1 of sensory preconditioning and advance our understanding of their differential/dissociable involvement in processing of the neutral S2-S1 association.

*Experiment 5A. Stimulating the LA increases inhibitory post-synaptic currents in the PRh.* We first recorded from visually identified layer 5 pyramidal neurons in the PRh in voltage clamp mode using a cesium-based internal solution. We chose layer 5 neurons because they have been shown to receive highly processed input from regions of the cortex, thalamus and, critically, the amygdala (Pikkarainen and Pitkänen, 2001). We recorded post-synaptic current responses at a range of membrane holding potentials from -83 mV to -7 mV (corrected for a +13 mV junction potential), while stimulating electrically within the LA (Fig. 6A). In all PRh neurons from which we recorded (*n* = 10), we found that LA stimulation produced a delayed post-synaptic response in the PRh neuron. The onset for these responses was between 5.0 and 11.4 ms post stimulus (average 8.1 ± 0.7 ms, *n* = 10) suggesting that these responses were not monosynaptic connections from LA inputs, but rather di-synaptic responses. The reversal potential for these currents (shown on the normalized I/V plot in Fig. 6A, right) was close to the expected reversal potential for chloride, calculated at -44 mV with adjustment for liquid junction potential, indicating that these responses were inhibitory post-synaptic currents (IPSCs). To confirm this, we recorded outward evoked currents at -13 mV and applied the GABA_A_ antagonist bicuculine (20 µM) to the perfusate and found this blocked the responses completely (Fig. 6B, average amplitude change was 101.9 ± 1.5 %, p < 0.05, *n* = 7). On blockade by bicuculine, we saw no inward current in any cell, suggesting that there is no direct evoked excitatory input to the layer 5 pyramidal neurons from the LA (*n* = 7). Upon washout of the bicuculine (Fig. 6B, *n* = 3) or with initial application to the evoked IPSC (*n* = 3), these currents were also completely blocked by the AMPA receptor-selective antagonist NBQX (10 µM, 101.7 ± 2.9% block, p < 0.05, *n* = 6). The implication of these results is that stimulation from the LA evoked an excitatory connection onto cortical interneurons which, in turn, provided the di-synaptic inhibitory input to the layer 5 PRh neurons.

**Figure 6.**
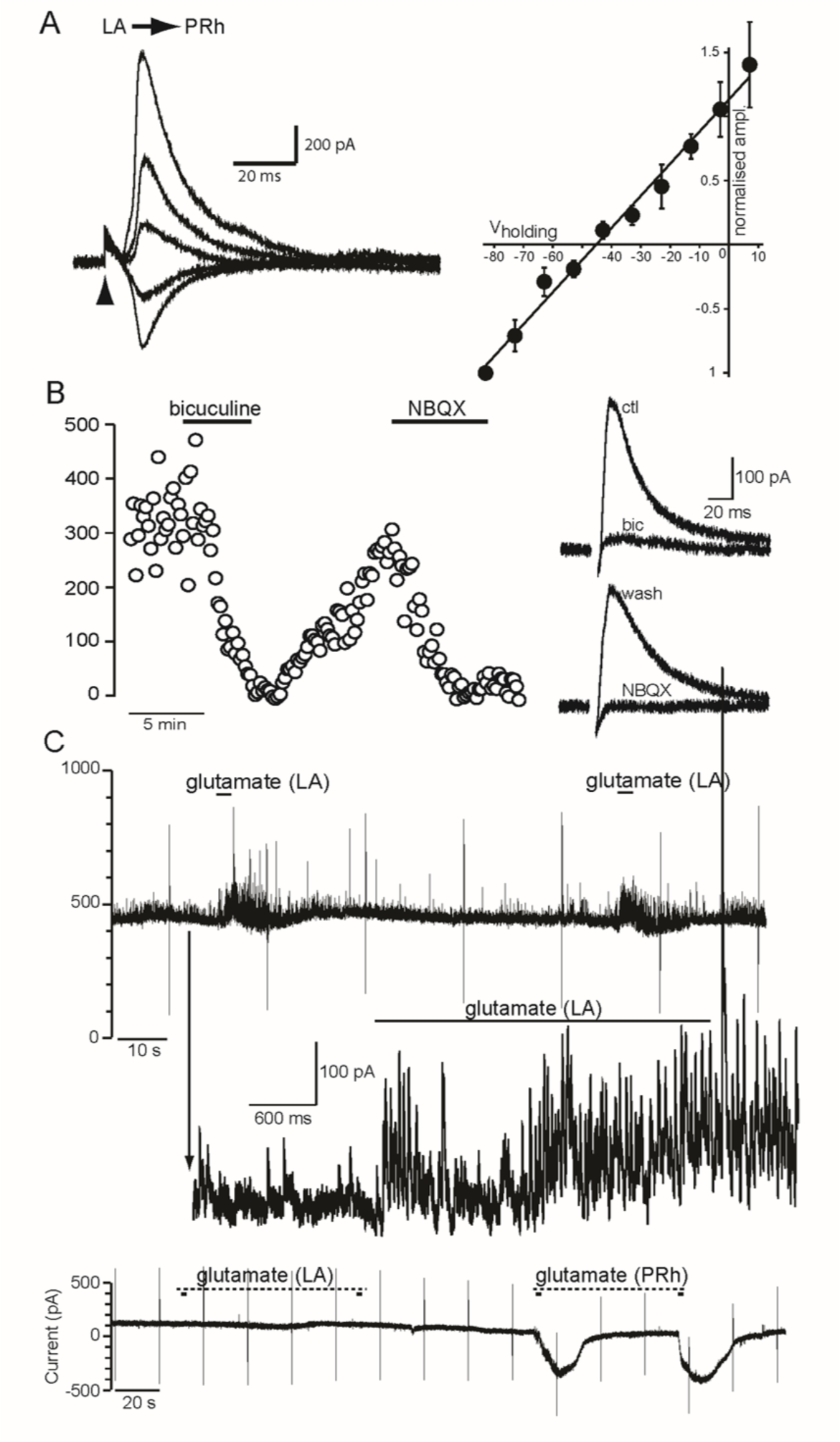
Synaptic responses of layer 5 PRh neurons following stimulation of the LA in Experiment 5A. **(A)** *Left:* Synaptic currents recorded from a layer 5 PRh neuron following electrical stimulation of LA. The stimulation timing is indicated with the arrowhead, and responses recorded at membrane potentials -3, -23, -43, -63 and -83 are shown (top to bottom overlaid). *Right:* Graph of the normalized current/voltage relation for responses recorded as shown in panel A, from membrane potential 7 to -83 mV and normalized to the amplitude of the -83 mV response (*n* = 7 cells). **(B)** *Left:* Amplitude plot showing peak evoked current amplitudes recorded at -13 mV membrane potential with the addition of bicuculine (20 µM) and NBQX (10 µM) to the perfusate. *Right:* Average evoked currents recorded in control (ctl) and in bicuculine (bic) and following washout of bicuculine (wash) and subsequent application of NBQX (NBQX) from this experiment. **(C)** The top chart recording shows an increase in outward synaptic currents recorded at -13 mV with pressure application of glutamate to layer 4/5 of the PRh. The middle trace shows the period before and during glutamate application on an expanded time axis. The lower trace shows a chart recording made at membrane potential -43 mV with glutamate applications made in the LA (first 2 applications) followed by two applications into an adjacent layer five region of PRh.

Electrical stimulation in a brain slice indiscriminately activates local cells within the field of the stimulator and axons passing through this region as fibers of passage. To confirm that responses seen in the PRh following electrical stimulation of the LA resulted from activation of LA neurons, we recorded from layer 5 neurons in PRh while applying the excitatory transmitter glutamate discretely into the LA using pressure injection from a 1-1.5 MΩ patch pipette. Application of glutamate in the LA would only affect responses in the PRh if it activated glutamate receptors in the region where it was applied, resulting in depolarization/firing of LA neurons and, thereby, LA-specific input to the PRh. Holding the perirhinal cells at -13mV (Fig. 6C), we found that glutamate puffs in the LA were immediately followed by increased IPSC frequency in 5 of 6 recorded neurons in the PRh. In some of these cases, the puffer pipette was relocated within the LA to achieve a response, suggesting that it is a sub-population of LA neurons that project to the PRh; and that the small volume of glutamate expelled in the LA achieved resulted in a discrete region of glutamate concentration that was rapidly diluted in the bath solution. To confirm that glutamate from the puff did not spread to activate interneurons in the PRh directly, we recorded from PRh neurons at the expected reversal potential for chloride (-47 mV) while puffing glutamate in the LA (*n* = 3). We saw no glutamatergic currents following puff application of glutamate in this configuration. Given the extent of the dendritic arborization of layer 5 cells within the cortical column, this suggests that glutamate was not spreading to the cortex in a concentration sufficient to activate neurons. By contrast, relocating the puffer pipette and applying glutamate within the PRh adjacent to the recorded cell produced large inward glutamate responses (Fig. 6C, *n* = 3), as expected. Thus, glutamate applied to the LA is unlikely to have directly activated interneurons in the PRh. Instead, we take these results to mean that glutamatergic excitation of LA neurons provided the glutamatergic drive for the di-synaptic inhibitory connection we had evoked electrically (though it should be noted that these experiments do not definitively rule out fibers of passage contributing to the electrically evoked responses).

*Experiment 5B. Stimulating the PRh increases inhibitory post-synaptic currents in the LA.* We next examined the capacity for inhibition from the PRh to LA. To do so, we recorded from LA neurons in voltage clamp mode with a cesium-based internal solution at a range of holding potentials, while stimulating electrically in layers 4 and 5 of PRh (Fig. 7A). We stimulated in layers 4 and 5 of the PRh because these regions exhibit clear projections to a range of cortical and subcortical areas, including the amygdala (McDonald et al., 1999; Pikkarainen & Pitkanen, 2001; Pitkanen et al., 2006; Shi & Cassell, 1999). This stimulation evoked delayed currents in LA neurons which were similar to those observed in PRh, with a comparable post stimulus onset (7.6 ± 0.4 ms). These currents also reversed close to the expected reversal potential for chloride (Fig. 7A, right) and were blocked by application of bicuculine (20 µM) or NBQX (10 µM; Fig. 7B; average block by bicuculine was 98.5 ± 0.5%, p < 0.01, *n* = 5, and average block by NBQX was 101.1 ± 2.5%, p < 0.01, n = 7). Moreover, discrete application of glutamate into the layer 4/5 region of PRh also resulted in increased IPSC frequency in 4 of 4 LA neurons that were tested (Fig. 7C). These results indicate that layer 4/5 PRh neurons provide excitatory input to interneurons connected to LA neurons, resulting in a di-synaptic inhibition of the LA neurons (though, as above, they do not definitively rule out a contribution from fibers passing through the PRh to the electrically stimulated responses).

**Figure 7.**
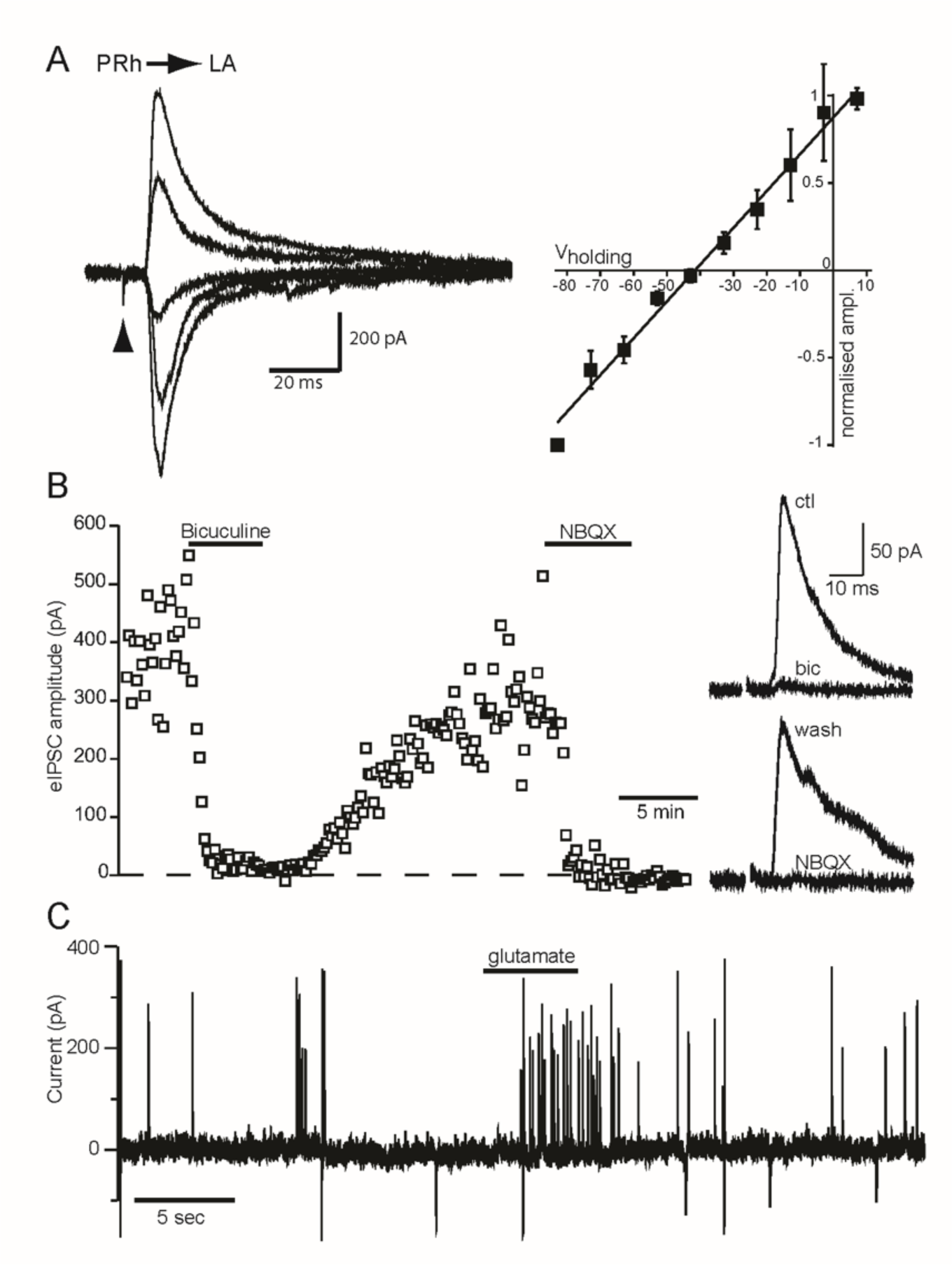
Synaptic responses of LA neurons following PRh stimulation in Experiment 5B. **(A)** *Left:* Synaptic currents recorded from a LA neuron following electrical stimulation of the layer 4/5 region of PRh. The stimulation timing is indicated with the arrowhead, and responses recorded at membrane potentials -3, -23, -43, -63 and -83 are shown (top to bottom overlaid). *Right:* Graph of the normalized current/voltage relation for responses recorded as shown in the left panel, from membrane potential 7 to -83 mV and normalized to the amplitude of the -83 mV response (*n* = 7 cells). **(B)** *Left:* Amplitude plot showing peak evoked current amplitudes recorded at -13 mV membrane potential with the addition of bicuculine (20 µM) and NBQX (10 µM) to the perfusate. *Right:* Average evoked currents recorded in control (ctl) and in bicuculine (bic) and following washout of bicuculine (wash) and subsequent application of NBQX (NBQX) from this experiment. **(C)** Chart recording showing an increase in outward synaptic currents recorded from a LA neuron at membrane potential -13 mV with pressure application of glutamate to layer 4/5 of the PRh.

In summary, these experiments show that stimulating the LA results in di-synaptic inhibition of pyramidal neurons in the PRh; and, conversely, that stimulating the PRh results in di-synaptic inhibition of pyramidal neurons in the LA. As such, they highlight a potential mechanism for the findings above showing that the PRh and BLA compete for processing of the neutral S2-S1 association in stage 1 of sensory preconditioning. When the S2-S1 pairings are novel, they activate neurons in the LA/BLA which then suppresses activity in the PRh: hence, processing of the early S2-S1 pairings requires activation of NMDA receptors in the BLA but not the PRh. By contrast, when the S2-S1 pairings have been repeated and the stimuli are familiar, they activate neurons in the PRh which then suppresses activity in the BLA: hence, processing of the later S2-S1 pairings requires activation of NMDA receptors in the PRh but not the BLA.

## Discussion

In this series of experiments, rats were exposed to pairings of two novel but affectively neutral stimuli (one auditory and the other visual, labelled S2 and S1) and, a day or so later, to pairings of S1 and shock. Rats froze when subsequently tested with the sensory preconditioned S2 and the conditioned S1. We and others have provided evidence that fear responses to S2 are not due to generalization from the conditioned S1 or any intrinsic ability of S1 to condition such responses to S2 (Holmes et al., 2013; Holmes and Westbrook, 2017; Kikas et al., 2021; Michalscheck et al., 2021; Parkes & Westbrook, 2010; Rizley & Rescorla, 1972; Wong et al., 2019). Rather they are associatively mediated: rats integrate the associations produced by the S2-S1 and the S1-shock pairings to generate fear responses to S2. The question of interest addressed here concerned the substrates of the S2-S1 association: specifically, whether danger, in the form of a shocked exposure to the context where the S2-S1 pairings had just occurred, affects the way that the association is consolidated in the PRh and/or BLA.

Experiments 1A and 1B showed that a shocked context exposure immediately after the session of S2-S1 pairings changes the way that the S2-S1 association is consolidated in the PRh and BLA. When this session was followed by a brief context alone exposure, the S2-S1 association was consolidated via protein synthesis-dependent changes in the PRh but not the BLA. By contrast, when this session was followed by a shocked context exposure, consolidation of the S2-S1 association ceased to require *de novo* protein synthesis in the PRh and, instead, required de novo protein synthesis in the BLA. That is, the shocked context exposure after sensory preconditioning had two consequences for consolidation of the just formed S2-S1 association: it engaged the BLA in support of this consolidation, and disengaged the PRh.

Subsequent experiments examined how danger in the form of the shocked context exposure engaged the BLA for consolidation of the S2-S1 association and disengaged the PRh. They tested the proposal that, during sensory preconditioning, the S2-S1 pairings generate two memory traces: one relating to the early novel pairings that is encoded in the BLA and another relating to the later familiar pairings that is encoded in the PRh. Events that occur after preconditioning then determine which of the two traces is consolidated to long-term memory. When rats are re-exposed to the context alone after preconditioning (i.e., if the context remains safe), the trace of the novel S2-S1 pairings in the BLA decays and the trace of the familiar S2-S1 pairings in the PRh is selected for consolidation. However, when rats are shocked in the context after preconditioning (i.e., the context becomes dangerous), the trace of the familiar S2-S1 pairings in the PRh is suppressed and the trace of the novel S2-S1 pairings in the BLA is selected for consolidation.

Experiments 2A and 2B confirmed the first part of this proposal: that the initially novel S2-S1 pairing is encoded in the BLA and not the PRh. Specifically, Experiment 2A demonstrated sensory preconditioning when rats are exposed to a single S2-S1 pairing in stage 1; and Experiment 2B then showed that the single S2-S1 pairing is encoded via activation of NMDAr in the BLA and not the PRh. Experiments 3A and 3B then confirmed the second part of the proposal: namely, that events which occur after the session of repeated S2-S1 pairings determine which of the BLA- or PRh-dependent traces is consolidated to the long-term memory system. Experiment 3A showed that NMDAr in the PRh are engaged by the repeated S2-S1 pairings in our standard sensory preconditioning protocol (repeated S2-S1 pairings), but that their involvement in sensory preconditioning is blocked when the preconditioning session is followed by a shocked context exposure. Experiment 3B showed that a shocked context exposure engages the BLA for consolidation of the S2-S1 association, but *only* if NMDAr in the BLA had been activated during the preconditioning session. Thus, the BLA (not PRh) supports the association produced by a single S2-S1 pairing and the PRh (not BLA) supports the association produced by multiple S2-S1 pairings. However, danger after the session containing multiple S2-S1 pairings cancels the involvement of the PRh and, instead, recruits the BLA to consolidate the sensory preconditioned association; but *only* if the BLA had been available to encode the initial pairing.

The final experiments extended our proposal by showing that danger after sensory preconditioning shifts the substrates of pre-extinction from the PRh to the BLA (Experiments 4A and 4B); and identified a potential mechanism by which the “either-or” roles of the PRh and BLA are achieved: stimulating the LA resulted in di-synaptic inhibition of pyramidal neurons in the PRh (Experiment 5A) and, conversely, stimulating the PRh resulted in di-synaptic inhibition of pyramidal neurons in the LA (Experiment 5B). More generally, the present findings extend our previous work which showed that the presence of danger *at the time of S2-S1 pairings* changes how the pairings are processed in the brain. When rats are exposed to the pairings in a safe and familiar context, the sensory preconditioned association is encoded through activation of NMDA receptors in the PRh, not the BLA (Holmes et al., 2013); and consolidated through molecular events in the PRh, not the BLA (Holmes et al., 2018). By contrast, when rats are exposed to S2-S1 pairings in a context that is equally familiar but dangerous (i.e., one in which rats are frightened because they had been shocked there previously), the sensory preconditioned association is encoded through activation of NMDA receptors in the BLA, not the PRh (Holmes et al., 2013); and consolidated through molecular events in the BLA, not the PRh (Holmes et al., 2018). These results were taken to imply that danger shifts encoding/consolidation of a sensory preconditioned association from the PRh to the BLA. However, the present findings show that this interpretation is incorrect. The initial S2-S1 pairings in sensory preconditioning are processed in the BLA (not the PRh) as these stimuli are novel and their consequences unknown. If the context is familiar and safe, the repeated S2-S1 pairings come to be processed in the PRh as they become increasingly familiar and their consequences (nothing) known. If, however, the context is dangerous, the repeated S2-S1 pairings continue to be processed in the BLA and are not processed in the PRh. That is, a dangerous context does *not* shift processing of the S2-S1 pairings from the PRh to the BLA. Rather, it *prevents* the shift in processing of the S2-S1 pairings from the BLA to the PRh: hence, sensory preconditioning in a dangerous context requires activation of NMDA receptors and molecular events in the BLA and not the PRh (these interpretations are schematized in Figure 8).

**Figure 8.**
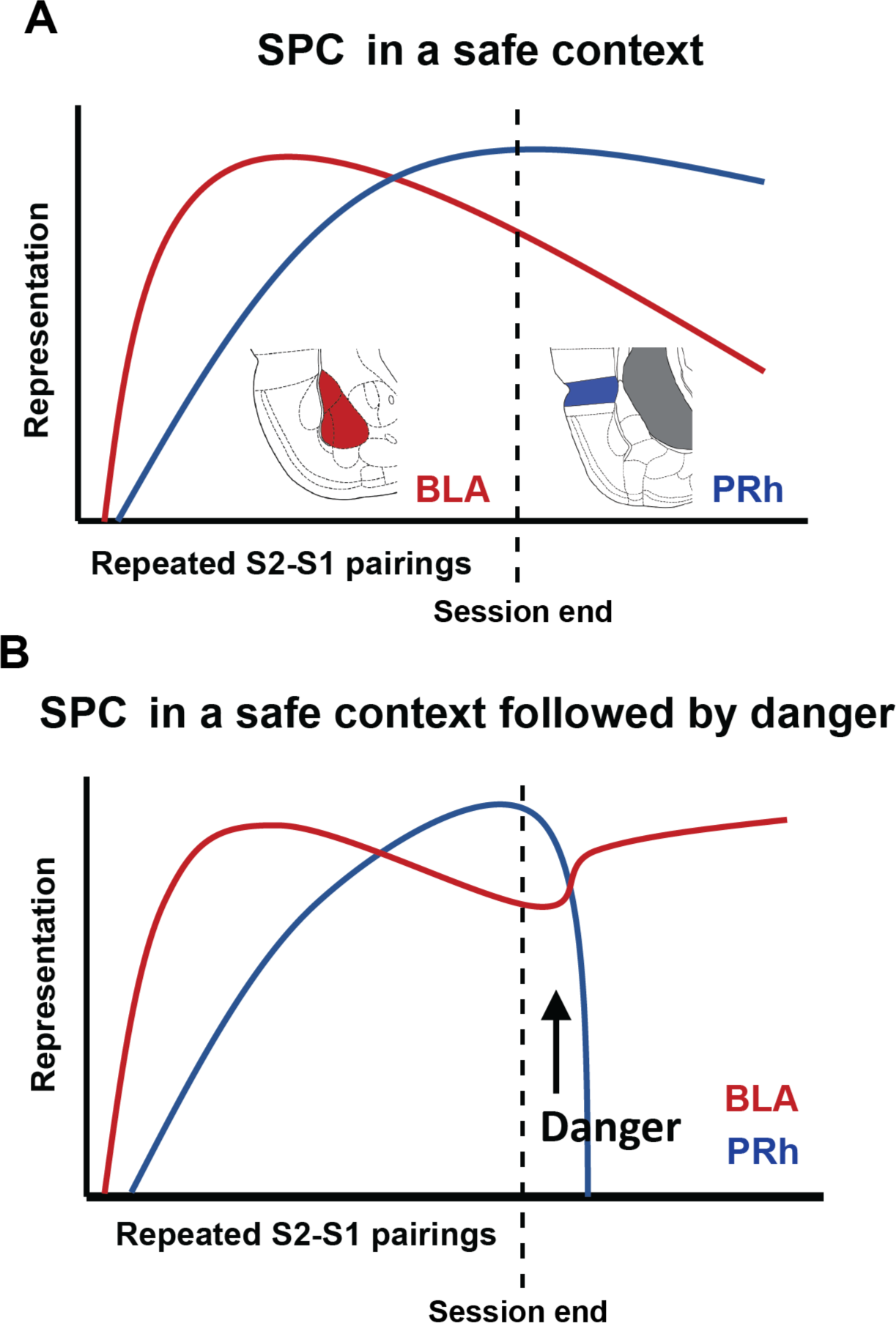
Schematic of putative memory traces in the PRh and BLA during and after sensory preconditioning (SPC) in a safe context (A) or a safe context followed by danger (B). A single S2-S1 pairing activates a memory trace in the BLA. Multiple S2-S1 pairings activate a memory trace in the PRh. When the sensory preconditioning session is followed by a context alone exposure, the trace in the BLA decays and the trace in the PRh is selected for consolidation (A). When the sensory preconditioning session is followed by danger, the trace in the PRh is suppressed and the trace in the BLA is selected for consolidation (B).

One additional point should be noted in relation to the present findings. While prolonged and severe stress disrupts memory in a range of protocols (McEwen and Sapolsky, 1995; Schwabe et al., 2009), acute and moderate stress has been shown to protect recently formed memories from forgetting (McGaugh, 2013) and to alter the substrates of successful memory in the amygdala. For example, Ritchey et al (2017) exposed people to a mild stressor (a cold-pressor test) immediately after they had viewed a series of pictures; and assessed the impact of the stressor on subsequent memory for those pictures. They found that successful memory in the stressed group was associated with higher levels of encoding-related activity in the amygdala: i.e., relative to remembered items in the non-stressed group, remembered items in the stressed group had elicited more activity in the amygdala during the encoding session. These findings are remarkably similar to those of the present study, where danger after sensory preconditioning engaged the BLA for consolidation of the innocuous S2-S1 association, *but only if the BLA had been available during the preconditioning session* (see also Shields et al., 2017). The similarity in findings across species and protocols reinforces the conclusion that post-encoding stress influences the substrates of memory in the amygdala and PRh, and that it does so by interacting with encoding-related activity in these regions. That is, post-encoding stress increases memory dependence on activity in the amygdala and decreases memory dependence on activity in the PRh.

In summary, the present study has shown that danger after sensory preconditioning disengages the PRh and engages the BLA to consolidate the new S2-S1 association; and identified these effects of danger with changes in processing of the S2-S1 association across the preconditioning session. As the initially novel stimuli become increasingly familiar, their processing shifts from the BLA to the PRh; but when danger occurs after the preconditioning session, the BLA-dependent trace of the novel S2-S1 pairings is consolidated and the PRh-dependent trace of the familiar S2-S1 pairings is suppressed. Future work will examine whether danger influences the consolidation of other types of memories by interacting with the processes by which they are encoded; and whether other types of emotional stimuli affect consolidation of recently acquired information in the same way.

## Acknowledgements

This work was supported by Australian Research Council (ARC) Future Fellowship to NMH (FT190100697), ARC Discovery Project Grants to NMH and RFW (DP200102969), and to RFW and SK (DP220103650), and an Australian Government Research Training Fellowship to OAQ.

